# Photon-HDF5: an open file format for single-molecule fluorescence experiments using photon-counting detectors

**DOI:** 10.1101/026484

**Authors:** A. Ingargiola, T. Laurence, R. Boutelle, S. Weiss, X. Michalet

## Abstract

We introduce Photon-HDF5, an open and efficient file format to simplify exchange and long term accessibility of data from single-molecule fluorescence experiments based on photon-counting detectors such as single-photon avalanche diode (SPAD), photomultiplier tube (PMT) or arrays of such detectors. The format is based on HDF5, a widely used platform- and language-independent hierarchical file format for which user-friendly viewers are available. Photon-HDF5 can store raw photon data (timestamp, channel number, etc…) from any acquisition hardware, but also setup and sample description, information on provenance, authorship and other metadata, and is flexible enough to include any kind of custom data.

The format specifications are hosted on a public website, which is open to contributions by the biophysics community. As an initial resource, the website provides code examples to read Photon-HDF5 files in several programming languages and a reference Python library (*phconvert*), to create new Photon-HDF5 files and convert several existing file formats into Photon-HDF5. To encourage adoption by the academic and commercial communities, all software is released under the MIT open source license.

## 1. Introduction

Single-molecule fluorescence techniques have facilitated ground-breaking developments in biophysics, as recently recognized by the Nobel committee (1). Techniques based on acquisition of long series of images obtained with ultra-sensitive cameras are the most popular among those, with numerous published software tools (2–4). A related set of single-molecule fluorescence techniques relying on photon-counting devices is used in studies of freely-diffusing single-molecules (5–7), studies of single-molecules in microfluidic mixers (8), optical or ABEL traps (9–11), or on immobilized molecules (12), where high temporal resolution (< 1 ms) is required. Among these techniques, single-molecule fluorescence resonant energy transfer (smFRET)(13), microsecond alternating laser excitation (µs-ALEX)(14), nanosecond ALEX (ns-ALEX)(15), also known as pulse interleaved excitation (PIE)(16), two-color coincident detection (TCCD)(17, 18), polarization anisotropy(19, 20), fluorescence correlation and cross-correlation spectroscopy (FCS, FCCS) (21, 22) or multi-parameter fluorescent detection (MFD)(23) are but a few in a constantly growing list.

Contrary to camera-based techniques, where most manufacturers provide a way to save data in a standardized format such as TIFF images, there is no such standardization for photon-counting hardware, where manufacturers of integrated systems have their own binary formats, and most systems are custom-built, including custom software generating oftentimes poorly documented binary files.

This situation creates a number of issues. First, it is rarely possible for an outsider (for instance a reviewer) to access the raw data of a published study without considerable coding work. Moreover, this assumes that the authors have provided a comprehensive description of their format. Worse, it is often difficult for different generations of students in a single laboratory to decipher data file formats created by their predecessors. In summary, the proliferation of file formats impedes interoperability between different software, creates barriers to data sharing among researchers and undermines results reproducibility and long-term preservation of data. Moreover, with the growing trend by funding agencies to request data sharing and documentation, having multiple file formats will inevitably result in significant duplication of efforts by each group.

To help address these issues, we developed Photon-HDF5(24), an open file format for single-molecule fluorescence experiments using photon-counting devices, including Time-Correlated Single-Photon Counting (TCSPC) data. The file format description comes accompanied with a reference Python library (*phconvert*)(25) to create Photon-HDF5 files from raw data or existing file formats, as well as examples(26) on how to read Photon-HDF5 files in Python, MATLAB and LabVIEW. Both format documentation and software are hosted on GitHub, a popular collaborative platform (27), with the aim of allowing the biophysics community to contribute to this project with comments, suggestions and code (28).

The article is organized as follows. In Section 2, we briefly review the current situation (characterized by a myriad of formats) and showcase a few successful examples of standardization in other scientific domains. In section 3, we examine requirements for an efficient, maintainable file format and introduce HDF5, a format on which Photon-HDF5 is based. We then describe the overall structure of Photon-HDF5 files and support software. In section 4, we illustrate the use of this format with a few scenarios, and in section 5, we conclude with a summary of Photon-HDF5 key features and with a call for the community to participate in shaping the evolution of the format.

## 2. Current situation

### 2.1 Photon-counting based experiments

Photon-counting based single-molecule fluorescence experiments can rapidly generate large amounts of data: in a typical experiment, count rates from a few kHz to several 10 kHz per detector and measurement durations of hours are not uncommon and, increasingly, large number of individual detectors are encountered (29–31). Since each photon’s information (time stamp, channel number, etc.) may occupy 64 or more bits per photon, most experiments store data in binary formats, which minimize disk space usage, at the expense of human readability. Because there is no natural or standardized structure for such a file format, each research group or company designing or putting together photon-counting instrumentation for single-molecule fluorescence experiments usually ends up developing its own custom binary file format.

In contrast with relatively well self-documenting text-based format, binary formats require detailed documentation of the file structure in order to be properly read (or written). Furthermore, extending a binary format to handle a new piece of information (such as modifications to the setup hardware) usually results in a new, incompatible format requiring specific handling and new, time-consuming code development. Since most researchers are, understandably, not inclined to invest significant resources in developing and documenting file formats, it is common to end up with a collection of poorly documented *ad hoc* binary formats within each individual research groups. The obvious risk is that, over the years, information about those file formats gets lost, effectively leading to data loss.

While vendors of photon-counting hardware and software have usually done a better job at designing and documenting file formats, their formats are nonetheless tightly linked to specific hardware and therefore, generally not suitable to store data from a competitor’s systems or from custom hardware used in individual laboratories. Additionally, although these formats can store some predefined metadata (such as for instance author, date of creation and comment), they are not generally extensible to accept additional metadata, necessary to describe a measurement in details.

The result of this situation is that it is difficult, if not impractical, for research groups not using identical hardware and software to exchange data to compare methods and results, and within each group, migration to new hardware or software is discouraged or results in incompatible data formats.

### 2.2 Examples of other formats

The need for standard file formats for domain-specific data sets is a common occurrence in many scientific fields. Most in the biophysics community will be familiar with the PDB file format for biomolecular structures maintained by the Protein Data Bank(32), or with the already mentioned TIFF format for images. The latter format, copyrighted by Adobe Systems, is only marginally adapted to the complexity of scientific data, but is used as a least common denominator export format by camera vendors in the absence of a better solution. The FITS format developed by the astronomy community(33) (and supported by some camera manufacturers) is a good example of a powerful alternative, although its design principles are dated and it has not been adopted by the biophysics or microscopy communities.

In the single-molecule community, a text-based file format (SMD) was recently proposed to store data from a variety of surface-immobilized, camera-based experiments(34). These measurements consist of series of images, from which time traces (time binned data) are extracted for individual emitters identified in the series. These time traces have relatively short duration (limited by fluorophore bleaching) and therefore have minimal space requirements. The proposed SMD format based on JavaScript Object Notation (JSON)(34) is therefore an appropriate choice for these structured data sets of relatively small size. On the other end, using SMD for timestamp-based experiments, would be inefficient (due to its text-based nature) and result in irreversible information loss (due to the time binning).

## 3. Proposal of a new file format

### 3.1 Design principles

The design of a standard file format for long-term data preservation and to facilitate interoperability and data sharing, presents several (often contradicting) requirements.

*Flexibility:* the format must be capable of efficiently storing a variety of numeric data types and strings to represent both raw data and all metadata associated with an experiment. Moreover, the structure must be general enough to accommodate different types of time stamp-based single-molecule experiments. For instance, it should support experiments using continuous wave or pulsed excitation, one or many detection channels, one or more excitation wavelengths, etc. Since it would be impossible to predict all possible future technical developments, it should be easily extensible and customizable to accommodate them.

*Ease of use:* in order to encourage adoption and facilitate long-term maintainability, the file structure and complexity should be kept as low as possible. It should be possible for a user to quickly write a reader for the file format in any programming language and operating system of choice. Furthermore, there should be tools to facilitate creation of data files that complies with the specifications.

*Small size:* while disk storage is increasingly inexpensive, the advantages of a compact file format should not be overlooked. Smaller files are easier to backup, archive online and share as well as generally faster to read. The format must therefore support smart compression allowing fast read and write, and partial loading of large data sets.

### 3.2 The parent HDF5 format

HDF5 is an existing open hierarchical file format satisfying all requirements listed previously. In the following, we provide a brief overview of the HDF5 format. Detailed information can be found on the HDF Group website(35).

HDF5 is a general-purpose binary format designed to store numerical data sets, widely used in industry and in several scientific communities. HDF5 development is led by the non-profit HDF Group, which also maintains open-source platform-independent libraries to read and write HDF5 files in multiple languages.

HDF5 has a hierarchical (tree-like) data structure similar to files and folders in a file system: groups are the equivalent of folders, whereas data fields (e.g. arrays) are the equivalent of files. The main group of a HDF5 file is called “root” (indicated by the slash character “/”) and can contain data fields or other groups. The HDF5 format is self-describing, meaning that a user does not need to know the hierarchical structure or the data types before reading the file. For example, in order to access a numerical array in a HDF5 file, the user only needs to specify its path: the array will be automatically loaded into memory using the appropriate data type. In other words, all the byte-level details (such as byte order and data type width) are transparently handled, greatly simplifying data reading and writing. Moreover, HDF5 transparently supports compression, allowing reducing file size while maintaining high reading speed without any added complexity for the user. Finally, each node in a HDF5 file (groups or data fields) can have any number of attributes, such as descriptions, which can be used to embed metadata to a dataset.

Libraries to operate on HDF5 files currently exist for the major programming languages used in scientific computing, such as C, Fortran, C++, Java, Python, R, MATLAB and LabVIEW. Furthermore, a free multi-platform graphical viewer and editor (HDFView (36)) allows users to quickly explore, modify or export content of HDF5 files. The availability of well-maintained open source libraries allows users to reliably manage HDF5 files without the need to deal with the underlying byte-level structure, effectively overcoming the drawbacks of binary formats highlighted in section 2.

HDF5 has been widely adopted in industry and in several scientific communities as the basis for specialized data formats. Because of its openness, popularity and long-term support, it represents a safe choice for long-term data preservation.

### 3.3 Photon-HDF5 file format

HDF5 is the foundation for the proposed format for photon-counting single-molecule fluorescence data, therefore called “Photon-HDF5” file format.

Photon-HDF5 is, essentially, a conventional structure to store time stamp-based single-molecule fluorescence data in HDF5 files (Fig. 1). Since Photon-HDF5 files are HDF5 files, the requirements of an open, efficient, well-documented and platform- and language-independent are automatically fulfilled.

**Figure 1:**
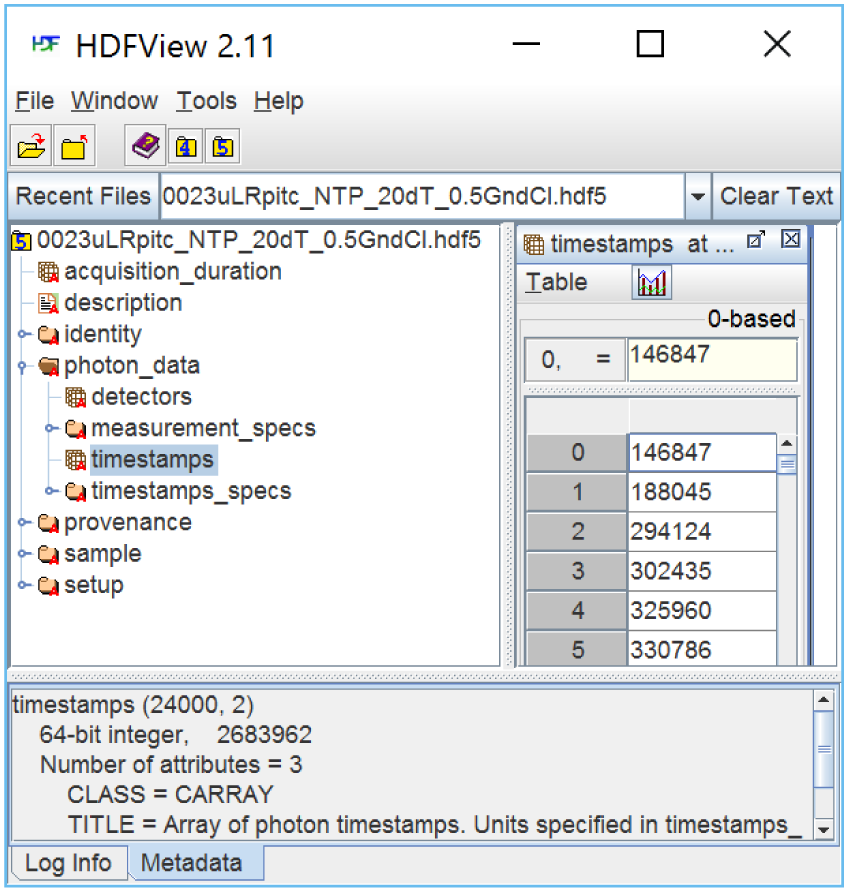
A Photon-HDF5 file as seen with the HDFView visualizer. In contrast with other binary formats, Photon-HDF5 files (or any other HDF5 file) can be examined with an open-source Java visualizer (HDFView). Note that the file structure (left pane) is immediately intelligible to the user, who can navigate it and select items for display or export. The time stamps array is selected in the left pane and its content displayed in the right pane. In the bottom pane, the TITLE attribute, among other attributes saved in the file, shows a description for the selected item (time stamps). Photon-HDF5 files include descriptions for every field, facilitating interactive file exploration.

We now briefly describe the main components of a Photon-HDF5 file. The interested reader is invited to visit the *photon-hdf5.org* website for further information and a complete description of the format and associated tools.

#### 3.3.1 Photon-HDF5 structure

The Photon-HDF5 file structure contains 5 main groups (*photon_data*, *setup*, *sample*, *identity* and *provenance*) and two root fields: a general description string (*description*) and the acquisition duration (*acquisition_duration*). The general structure is illustrated in Fig. 2 (see also Fig. SM-1 and SM-2), while Fig. 1 shows a Photon-HDF5 file as it appears when opened with the HDFView visualizer.

**Figure 2.**
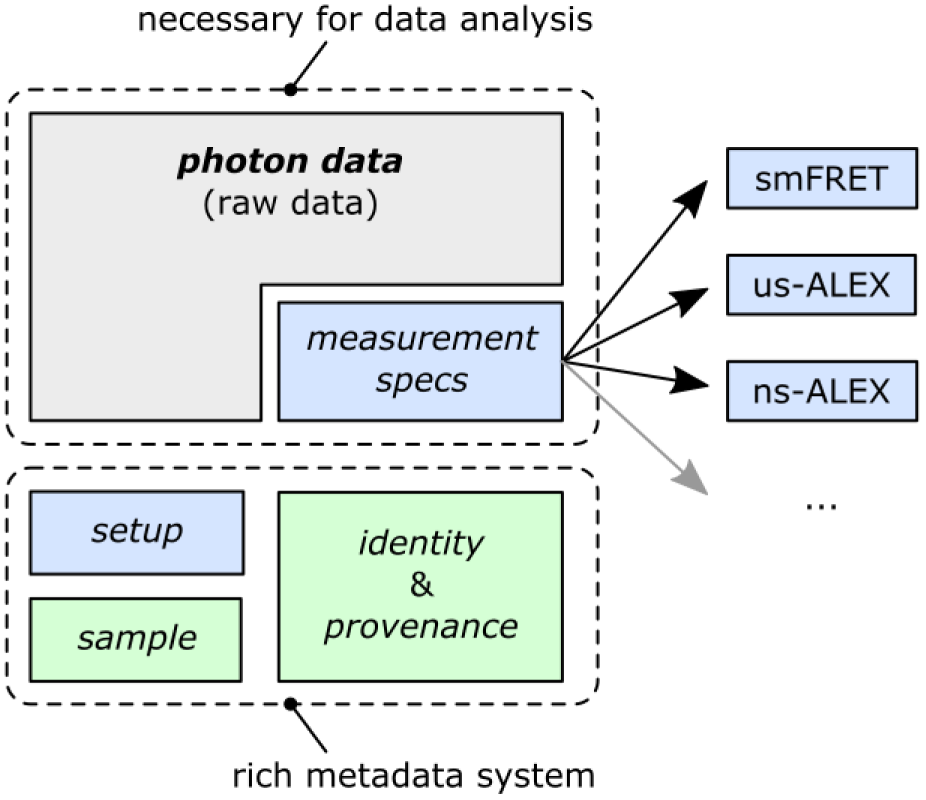
Photon-HDF5 file structure. The core *photon_data* group contains the raw data. A file containing only raw data is still a valid Photon-HDF5 file but it cannot be analyzed without additional information on what the data represents. The optional (but recommended) *measurement_specs* group identifies the type of measurement and specifies the associated metadata (i.e. role of each detector, measurement parameters, etc). When *measurement_spec* is present, the user has all information needed to analyze the data. The other groups enhance the data file legibility, providing important measurement metadata. *setup* contains information on the system used for acquisition (e.g. excitation wavelengths, laser power(s), CW or pulsed excitation, etc). *sample* contains sample information (e.g. dyes, buffer, etc…). The last two groups *identity* and *provenance* provide author and file information.

##### a. The *photon_data* group

As visible in Fig. 1 (and detailed in Fig. SM-1), the *photon_data* group contains the raw data (*timestamps*, *detectors* and *nanotimes* arrays), as well as information needed to analyze it (in the *measurement_specs* subgroup). All arrays in *photon_data* have the same length, equal to the number of recorded photons. Information corresponding to the i^th^ photon (its macrotime stamp, detector identity, and when applicable, the microtime measured by TCSPC hardware, or nanotime) is stored the i^th^ value of each of these arrays.

Except for *timestamps*, the other arrays are present in a file only when needed, i.e. the *detectors* array is present only if there is more than one detector and the *nanotimes* array is present only if TCSPC timing data is recorded.

Inside *photon_data*, the *measurement_specs* group contains the measurement type and a type-dependent set of metadata fields. The currently supported measurement types are listed in Table SM-1. Each measurement type (specified in *measurement_type*) requires a particular set of fields which completely describe the experiment. For example, a single-laser single-molecule FRET data file will have a “smFRET” measurement type, and will include two numeric fields called *spectral_ch1* and *spectral_ch2* containing the donor and acceptor detector IDs.

Defining a new measurement type involves choosing a “name” and a set of required fields in the *measurements_specs* group. Defining new measurement types allows gradually extending the file format in order to support new measurement types, without compromising the legibility of already existing files.

Data contained in *measurement_specs* group provides all the necessary information needed to interpret raw experimental data. Files not including a *measurement_specs* group are still valid Photon-HDF5 files, but a user cannot analyze their content without knowing additional measurement specifications. Such an option is provided only to store incomplete or debugging data and is not recommended for data that need to be shared. For new measurement types, users should submit new measurement_specs definitions to the community for integration in new releases of the photon-hdf5 format specifications(37).

##### b. The *setup* group

The *setup* group stores setup properties such as laser excitation wavelengths and powers, the number of detected spectral bands (e.g. 2-colors or 3-colors), the number of detected polarization states (e.g. none = 1 and 2 polarizations = 2) and the number of “split channels” (i.e. identical spectral or polarization channels split by non-polarizing beam-splitters).

##### c. Metadata groups

In addition to the previous groups, a Photon-HDF5 file may contain three other groups (*sample*, *identity* and *provenance*) which provide additional optional metadata (see Fig. 2, & SM-2).

The *sample* group contains a description of the sample (*sample_name*), the buffer (*buffer_name*) and the fluorophore names (*dye_names*). Storing this basic sample information with the data simplifies preservation of experimental details.

The *identity* group contains identification metadata such as dataset *author* (and its affiliation), Photon-HDF5 format version, software used to save the file (and its version), and optional dataset identifiers such as DOI or URL. This information is important when sharing datasets online, in order to keep track of authorship and provide proper credit.

As discussed in Section 4, a Photon-HDF5 file might be created by converting an original data file saved by an acquisition (or simulation) software outputting some custom file format. The *provenance* group is the place to store this information, such as the name and creation date of the original file, and the software used to save it.

#### 3.3.2 Additional file format features

##### a. Field data types

Leveraging the self-describing nature of HDF5, instead of prescribing specific data types, Photon-HDF5 specifications only indicate whether a field is a scalar, an array or a string and whether the data type is integer or floating point. For examples, the *detectors* array (array of detector IDs) typically uses an unsigned 8-bit integer type, which is sufficient for the majority of setups. However, when using SPAD arrays with more than 256 pixels, a 16- or 32-bit integer data type can be used without any modification to the file format.

##### b. Custom user groups

For maximum flexibility, users can store arbitrary amounts of additional (custom) data in any position within the Photon-HDF5 hierarchy. To avoid incompatibilities with future versions of the format, it is required that custom data be stored inside a group (or subgroup) named *user*.

##### c. Future Photon-HDF5 versions

In order to guarantee long-term accessibility to Photon-HDF5 files, future versions of the file format software will be backward-compatible. Specifically, previously defined fields will never be renamed or have their meaning modified. This ensures that software written to read an old version will be still capable to correctly read a newer version, although it will not be able to interpret newer fields or measurement types.

#### 3.3.3 Reading and Writing Photon-HDF5

Reading Photon-HDF5 files does not differ from reading any other HDF5 file and involves using the HDF5 library for the language of choice. A specific advantage is that field names (and their meaning) are defined in the specifications (SM.1). Examples of how to read Photon-HDF5 files in several languages are provided in SM.2 and reference (26) (currently for Python, MATLAB and LabVIEW). Additional discussion on reading Photon-HDF5 files can be found in the “Reading Photon-HDF5 files” section of the reference documentation (38).

To create Photon-HDF5 files, users can convert existing files or save them directly from suitably modified acquisition software. Conversion is generally the simplest approach and, when using acquisition software whose source code is unavailable, the only possible one. To simplify saving and converting Photon-HDF5 files, we maintain *phconvert*, an open-source Python library serving as reference implementation for the Photon-HDF5 format(25). *phconvert* includes a browser-based interface using Jupyter Notebooks(39, 40) to convert supported file formats into Photon-HDF5. The formats currently supported are HT3 (from PicoQuant TCSPC hardware), SPC/SET (from Becker & Hickl TCSPC hardware) as well as SM (a legacy file format used in the Weiss Lab). We provide an online demonstration service to run these notebooks and convert these formats to Photon-HDF5 without software installation on the user’s computer(41).

Additionally, the phconvert library can be used in any Python program to easily create Photon-HDF5 files from scratch (see SM.3). While Photon-HDF5 files can be created using only a HDF5 library (e.g. pytables or h5py), one advantage of using phconvert is that files are validated and guaranteed to conform to the specifications. Phconvert, in fact, checks that all fields have correct names and types, and adds a description to each field (see Fig. 1 for an example). Furthermore, phconvert automatically computes and fills several fields (e.g. file creation and modification dates, format and software names and versions, acquisition duration, etc…) reducing the amount of metadata the user needs to provide. Programs written in languages different from Python, can benefit from the same advantages of using phconvert for creating Photon-HDF5 files by calling a simple script called phforge(42) whose use is described in SM.3.2.

Other currently unsupported formats can be converted to Photon-HDF5 simply writing a Python function to load the data in memory and using phconvert to save the data to Photon-HDF5. Taking the existing loader functions as examples, this task is relatively easy, even for inexperienced Python programmers. We encourage interested users to contribute to phconvert so that out-of-the-box support for conversion of the largest number of formats can be provided. For a discussion on saving Photon-HDF5 from Python or other languages see SM.3 and the reference documentation (43).

## 4. Scenarios of Photon-HDF5 usage

In this section, we illustrate a few use cases where employing the Photon-HDF5 format may lead to significant advantages.

Researchers performing (or interested in) photon-counting based single-molecule experiments can be classified in the following three categories. Users using:

a. Commercial acquisition hardware and software (Alice)
b. Custom acquisition hardware and software (Bob)
c. Mixture of a and b (Charlie)

In each case, Alice, Bob and Charlie use one or more custom or proprietary binary formats to store data from their experiments, and will use a custom or commercial software to analyze their data. We now examine three typical situations encountered by researchers.

### Example 1 (Publication)

Alice wants to publish an article including her data so that as many researchers as possible can look at it (including the reviewers, who may not, however, have time to process it and might just be curious to take a peek at it).

The Photon-HDF5 format and HDFViewer will do the job for the reviewer and researchers using a different system, while the original format will allow only readers having the same commercial software to have a look at the data. If she uses one of the two most common commercial formats, she can use phconvert to create the Photon-HDF5 version. If not, writing code to create a Photon-HDF5 version should be simple thanks to the examples provided. Moreover, her code could be added to phconvert for the benefit of the entire community.

### Example 2 (Collaboration)

Alice and Bob want to collaborate on a project, Alice doing the experiments and Bob performing some custom analysis with his software.

Since Bob doesn’t use Alice’s system, he would have to decipher the proprietary file format description of the vendor and extract the relevant data for his analysis. By asking Alice to first convert her file to the Photon-HDF5 format with phconvert, he will benefit from file size reduction, simple access to the data, and by incorporating the necessary code in his software, he will gain access to a variety of other files generated by other users of Photon-HDF5.

### Example 3 (Data Management)

Charlie works with a student running some experiments on a commercial system, which he wants to compare to similar experiments he is carrying out on a new setup he has built with custom hardware and software.

Since Charlie needs to save new data, he needs a new format. But he also needs to read data generated by the student. The least effort consists in using a format which is efficient, well-documented and supported and in which his student’s files can be easily converted. Photon-HDF5 naturally fits the bill, and additionally provides him with the same benefits enjoyed by Alice and Bob for publication and sharing.

### Summary

While these scenarios do not directly involve manufacturers of single-photon counting hardware and their associated software, it is clear that manufacturers should take some interest in an effort such as Photon-HDF5. While it is not expected that they would abandon their proprietary format, their contribution to the definition of future versions of the format or to conversion tools is welcome and encouraged, as is that of anyone interested in the research community.

## 5. Conclusions & Perspectives

We have developed the Photon-HDF5 file format as a hardware-agnostic format for single-molecule fluorescence data based on photon-counting detectors. Photon-HDF5 is easy to use, customize and extend, in addition to inheriting all the advantages of the HDF5 format (open standard, self-described and efficient).

The format has been designed both for interoperability between different analysis software and long-term data archival. For this purpose, it includes a rich set of metadata to make data files self-contained. Photon-HDF5 facilitates data sharing between research groups by reducing the efforts needed to decode the wide variety of binary formats used across laboratories.

While we have only discussed standard single-spot confocal experiments in the text, the format supports and is currently used for multi-spot experiments using multi-pixel SPAD arrays(24). Further extensions to accommodate future developments or new measurement types can be naturally incorporated.

Currently, the open source FRETBursts software package for smFRET burst analysis supports reading Photon-HDF5 files(44). Therefore, converting smFRET data to Photon-HDF5 already enables researchers to analyze a variety of data with this software. The PyBroMo simulator for smFRET experiments generates simulated smFRET data in Photon-HDF5 format(45), which can be analyzed by FRETBursts or other software supporting Photon-HDF5. Additional software packages supporting Photon-HDF5 files are expected to be released in the near future and links to their websites will be added to the photon-hdf5.org site.

The impact of Photon-HDF5 format will be determined by the extent of its adoption in the single-molecule community and its support by scientific publishers. To maximize the involvement of third parties, we embraced an open development model for both the format specification and the support software. Users are encouraged to send feedback (e.g. on perceived limitations or to add support for new measurement types) or other contributions. To facilitate the process, we include guidelines for contributors in the Photon-HDF5 reference documentation(24). Finally, by adopting an open consensus-based governance, we aim to empower the single-molecule community to determine the future directions of Photon-HDF5 format in order to become a standard widely-useful tool for data sharing and preservation.

## Author Contributions

AI designed the file format, implemented the supporting software, wrote the reference documentation, websites and manuscript; TL designed the file format, implemented LabVIEW reading examples, wrote the manuscript; RB implemented the MATLAB code for reading and writing Photon-HDF5 files; SW wrote the manuscript; XM designed the file format, implemented LabVIEW reading and writing examples, wrote the manuscript.

## Acknowledgements

We thank Sangyoon Chung and Dr. Eitan Lerner for providing experimental data files. This work was supported in part by NIH Grant R01-GM95904 and by DOE Grant DE-FC02-02ER63421-00. Dr. Weiss discloses equity in Nesher Technologies and intellectual property used in the research reported here. The work at UCLA was conducted in Dr. Weiss's Laboratory. Dr. Laurence's work was performed under the auspices of the U.S. Department of Energy by Lawrence Livermore National Laboratory under Contract DE-AC52-07NA27344.

## SUPPORTING MATERIAL

### SM1. Photon-HDF5 files examples

In this section, we describe a few examples of Photon-HDF5 files corresponding to different types of measurement, focusing for brevity on the mandatory fields in */photon_data*. The fields here shown, as well as all the other fields in the remaining groups (*/setup*, */sampl*e, */identity* and */provenance*) can be found in the section “Photon-HDF5 format definition” of the reference documentation(1).

Photon-HDF5 files can be distinguished from other HDF5 files by the presence of two root node attributes *format_name* and *format_version*: the former contains the string “Photon-HDF5” and the latter is a string identifying the Photon-HDF5 version (currently “0.4”). The measurement type is identified by reading the string in */photon_data/measurement_specs/measurement_type*, whose currently supported values are reported in Table SM-1.

#### SM1.1 μs-ALEX

In μs-ALEX smFRET data files, the following data fields are present (their definitions can be found in the reference manual(1)):

1. Timestamps (array): /photon_data/timestamps
2. Timestamps unit (scalar): /photon_data/timestamps_specs/timestamps_unit
3. Detectors array (array): /photon_data/detectors
4. Measurement type (string): /photon_data/measurement_specs/measurement_type (“smFRET-usALEX”)

Additionally, the /photon_data/measurement_specs group contains the fields indicated in Table SM-1, row smFRET-usALEX.

#### SM1.2 ns-ALEX

In ns-ALEX smFRET (also known as PIE) data files, the following data fields are present:

1. Timestamps (array): /photon_data/timestamps
2. Timestamps unit (scalar): /photon_data/timestamps_specs/timestamps_unit
3. Detectors array (array): /photon_data/detectors
4. TCSPC nanotimes (array): /photon_data/nanotimes
5. TCSPC bin width (scalar): /photon_data/nanotimes_specs/tcspc_unit
6. TCSPC number of bins (scalar): /photon_data/nanotimes_specs/tcspc_num_bins
7. Measurement type (string): /photon_data/measurement_specs/measurement_type (“smFRET-nsALEX”)

Additionally, the /photon_data/measurement_specs group contains the fields indicated in Table SM-1, row smFRET-nsALEX. Definitions for all these fields can be found in the “Photon-HDF5 format definition” section of the reference manual(1).

**Table SM-1:** List of currently supported measurement types. Additional types can be added in the future based on user demand. The first column represents the string identifying the measurement type that is located at */photon_data/measurement_specs/measurement_type* in the Photon-HDF5 file. The second column is a brief description of the measurement type. The third column lists the required metadata fields in */photon_data/measurement_specs/* for each type of measurement.

| **Measurement type** | **Description** | **Required fields in measurement_specs** |
| --- | --- | --- |
| *smFRET* | 2-colors smFRET with single wavelength excitation | • `detectors_specs/spectral_ch1`<br>• `detectors_specs/spectral_ch2` |
| *smFRET-usALEX* | 2-colors smFRET with 2 alternating CW excitation lasers | • `detectors_specs/spectral_ch1`<br>• `detectors_specs/spectral_ch2`<br>• `alex_period`<br>• `alex_offset`<br>• `alex_excitation_period1 (optional)`<br>• `alex_excitation_period2 (optional)` |
| *smFRET-usALEX-3c* | 3-colors smFRET with 3 alternating CW excitation lasers | • `detectors_specs/spectral_ch1`<br>• `detectors_specs/spectral_ch2`<br>• `detectors_specs/spectral_ch3`<br>• `alex_period`<br>• `alex_offset`<br>• `alex_excitation_period1`<br>• `alex_excitation_period2`<br>• `alex_excitation_period3` |
| *smFRET-nsALEX* | 2-colors TCSPC-based smFRET with 2 alternating pulsed excitation lasers (also known as PIE) | • `detectors_specs/spectral_ch1`<br>• `detectors_specs/spectral_ch2`<br>• `laser_repetition_rate`<br>• `alex_excitation_period1 (optional)`<br>• `alex_excitation_period2 (optional)` |

**Figure SM-1:**
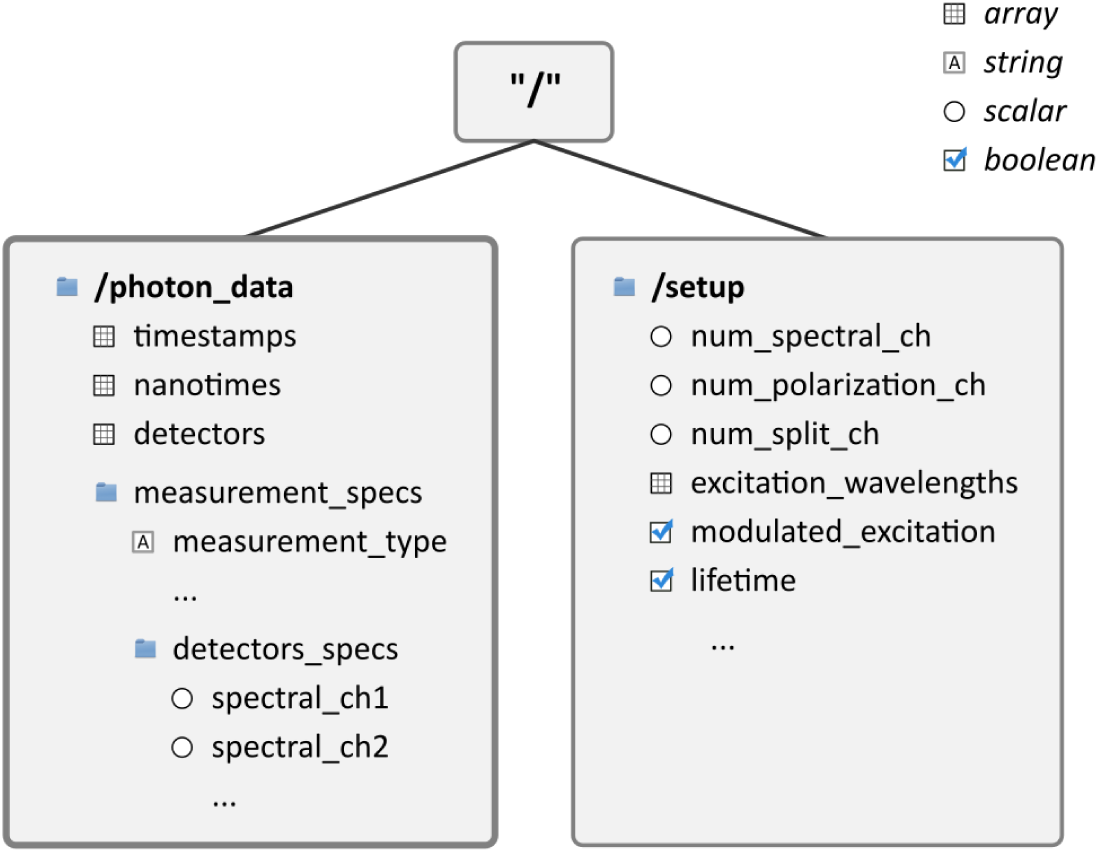
Partial diagram description of the Photon-HDF5 layout comprising the *photon_data* and *setup* groups, which contains the acquisition data and all the information needed for data processing. The symbol preceding each field indicates the data type of that field (e.g. array, string, scalar or boolean), as described in the legend.

**Figure SM-2:**
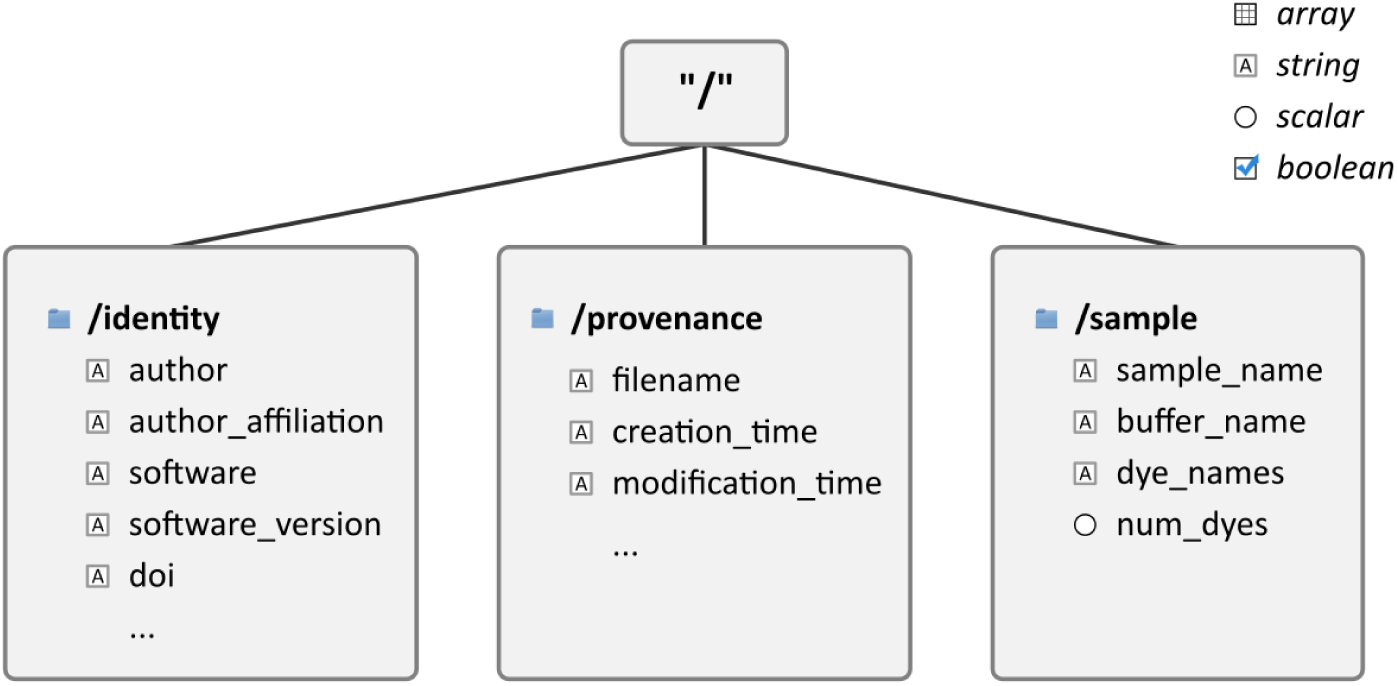
Partial diagram description of the Photon-HDF5 layout showing the *sample*, *identity* and *provenance* groups. These groups are not strictly necessary to analyze the data, but are important for long-term preservation, reproducibility and proper credit/citation when the file is shared.

### SM2. Reading Photon-HDF5 Files

In this section, we show basic examples of how to read a Photon-HDF5 file containing μs-ALEX data in Python (SM2.1), MATLAB (SM2.2) and LabVIEW (SM2.3). These and other examples of reading Photon-HDF5 files are provided online (2). Additional information on reading Photon-HDF5 files can be found in the section “Reading Photon-HDF5 files” of the reference documentation(3).

#### SM2.1 Reading in Python

Using Python and the pytables, we start by opening the data file and getting a handle to the photon-data group:

~~~
h5file = tables.open_file(filename)
photon_data = h5file.root.photon_data
~~~

Next, to read the timestamps and detectors array and the timestamps units, we use:

~~~
timestamps = photon_data.timestamps.read()
timestamps_unit = photon_data.timestamps_specs.timestamps_unit.read()
detectors = photon_data.detectors.read()
~~~

To retrieve donor and acceptor detector ID and other μs-ALEX specific fields:

~~~
donor_ch = photon_data.measurement_specs.detectors_specs.spectral_ch1.read()
acceptor_ch = photon_data.measurement_specs.detectors_specs.spectral_ch2.read()
alex_period = photon_data.measurement_specs.alex_period.read()
offset = photon_data.measurement_specs.alex_offset.read()
donor_period = photon_data.measurement_specs.alex_excitation_period1.read()
acceptor_period = photon_data.measurement_specs.alex_excitation_period2.read()
~~~

Finally, some summary information can be printed as follow:

~~~
print('Number of photons: %d' % timestamps.size)
print('Timestamps unit: %.2e seconds' % timestamps_unit)
print('Detectors:  %s' % np.unique(detectors))
print('Donor CH: %d Acceptor CH: %d' % (donor_ch, acceptor_ch))
print('ALEX period: %4d \nOffset: %4d \nDonor period: %s \nAcceptor period: %s' % (alex_period, offset, donor_period, acceptor_period))
~~~

This and other Python examples (using both pytables and h5py) can be found at:

> https://github.com/Photon-HDF5/photon_hdf5_reading_examples/tree/master/python

#### SM2.2 Reading in MATLAB

The following example works in MATLAB 2013a or later. For earlier versions of MATLAB (7.3 or later), the user may use the low-level HDF5 functions (which are however more complex).

To load timestamps and detectors arrays and timestamps unit, we execute:

~~~
timestamps = h5read(filename, '/photon_data/timestamps');
timestamps_unit = h5read(filename, '/photon_data/timestamps_specs/timestamps_unit');
detectors = h5read(filename, '/photon_data/detectors');
~~~

Next, to retrieve donor and acceptor detector ID and other μs-ALEX specific fields:

~~~
donor_ch = h5read(filename, '/photon_data/measurement_specs/detectors_specs/spectral_ch1');
acceptor_ch = h5read(filename, '/photon_data/measurement_specs/detectors_specs/spectral_ch2');
alex_period = h5read(filename, '/photon_data/measurement_specs/alex_period');
offset = h5read(filename, '/photon_data/measurement_specs/alex_offset');
donor_period = h5read(filename, '/photon_data/measurement_specs/alex_excitation_period1');
acceptor_period = h5read(filename, '/photon_data/measurement_specs/alex_excitation_period2');
~~~

And to print some summary information:

~~~
fprintf('Number of photons: %d\n', size(timestamps));
fprintf('Timestamps unit: %.2e seconds\n', timestamps_unit);
fprintf('Detectors: %s\n', unique(detectors));
fprintf('Donor CH: %d Acceptor CH: %d\n', [donor_ch; acceptor_ch]);
fprintf('ALEX period: %d \nOffset: %d \nDonor period: %d, %d Acceptor period: %d, %d\n',… [alex_period; offset; donor_period; acceptor_period]);
~~~

The full example can be found at:

> https://github.com/Photon-HDF5/photon_hdf5_reading_examples/tree/master/matlab

#### SM2.3 Reading in LabVIEW

Figure SM-3 shows a snapshot of one of the two LabVIEW codes provided as examples at

> https://github.com/Photon-HDF5/photon_hdf5_reading_examples/tree/master/labview

The code is saved in LabVIEW 2010 and should be compatible with later versions of LabVIEW. The code uses the h5labview wrapper for the HDF5 library. This requires a two-step installation. First, installation of the HDF5 library available at https://www.hdfgroup.org/ftp/HDF5/releases/hdf5-1.8.14/bin/windows/. At the time of this writing, the current tested version of the library is 1.8.14. Second, installation of h5labview (available at http://sourceforge.net/projects/h5labview/files/) using VPI Manager, a free utility to install LabVIEW packages. The examples have been generated using version 2.11.2.137 of h5labview2, but should work with later versions as well.

The only user input is the path of the file to be read (Path parameter on the front panel - not shown on Fig. SM-3). Pressing the “Run” button will load the file and display some basic information about its content, the raw data (timestamp and detector arrays) as well as represent the time stamp histogram modulo the alternation period and the donor and acceptor emission periods as they were defined when saving the file.

**Figure SM-3:**
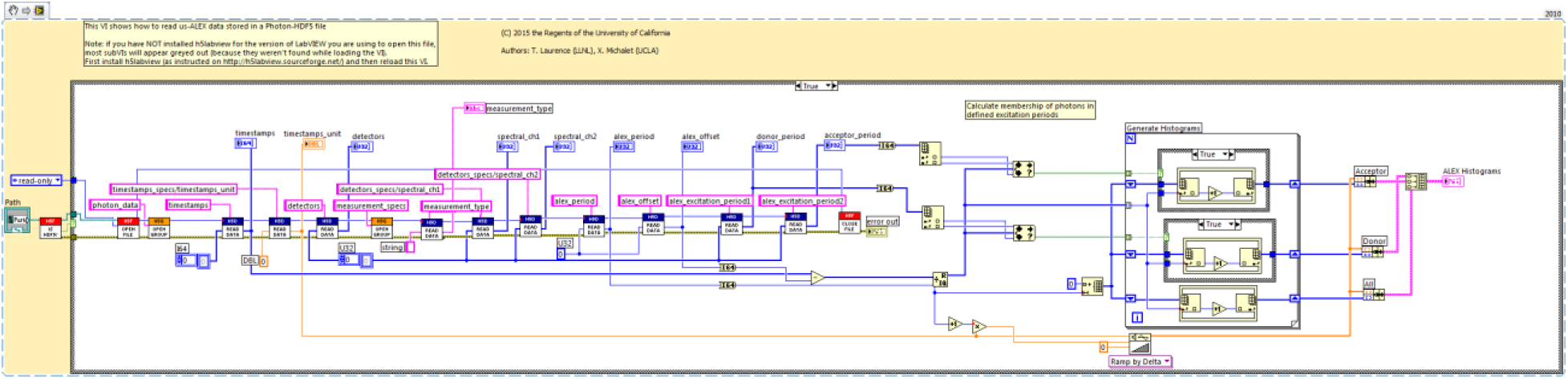
Example LabVIEW 2010 code to read a Photon-HDF5 file corresponding to a μs-ALEX measurement.

### SM3. Writing Photon-HDF5 Files

In the following sections, we continue the discussion of section 3.3.3 in the main text, on converting or saving Photon-HDF5 files.

As mentioned before, supported file formats can be directly converted using one of the Jupyter notebooks included in phconvert. For unsupported file formats, a user can write a Python function to load the data, and use *phconvert* to save it into the Photon-HDF5 file.

Users may want to implement the ability to save Photon-HDF5 files in software for data-acquisition or simulators. For Python programs the direct solution is calling phconvert to save the file as illustrated in SM-3.1. As mentioned before, using phconvert greatly simplifies writing Photon-HDF5 files while ensuring compatibility with the Photon-HDF5 specifications. For non-Python programs which cannot easily call phconvert, saving Photon-HDF5 files is equally easy (and reliable) using the phforge script(4) as described in SM-3.2. Under the hood, phforge calls the phconvert library and therefore creates files which are guaranteed to be valid Photon-HDF5 files.

#### SM3.1. Saving Photon-HDF5 from Python programs: phconvert

In this section, we provide an example of how to write a Photon-HDF5 in Python with phconvert. This example illustrates how to build the data structure needed to save a Photon-HDF5 file. This example can be also used as a reference by users willing to implement a loader function for an additional format.

The logic is the following. Each group in a Photon-HDF5 file is represented in memory by a Python dictionary: a key-value pair (in the dictionary) represents a field name and its content respectively (in the Photon-HDF5 file). For example, the root group dictionary contains an item whose key is 'description' (i.e. the HDF5 field name) and whose value is a string (i.e. the HDF5 field content).

Following this logic, the two mandatory groups of a Photon-HDF5 file (*photon_data* and *setup*) can be defined as follows:

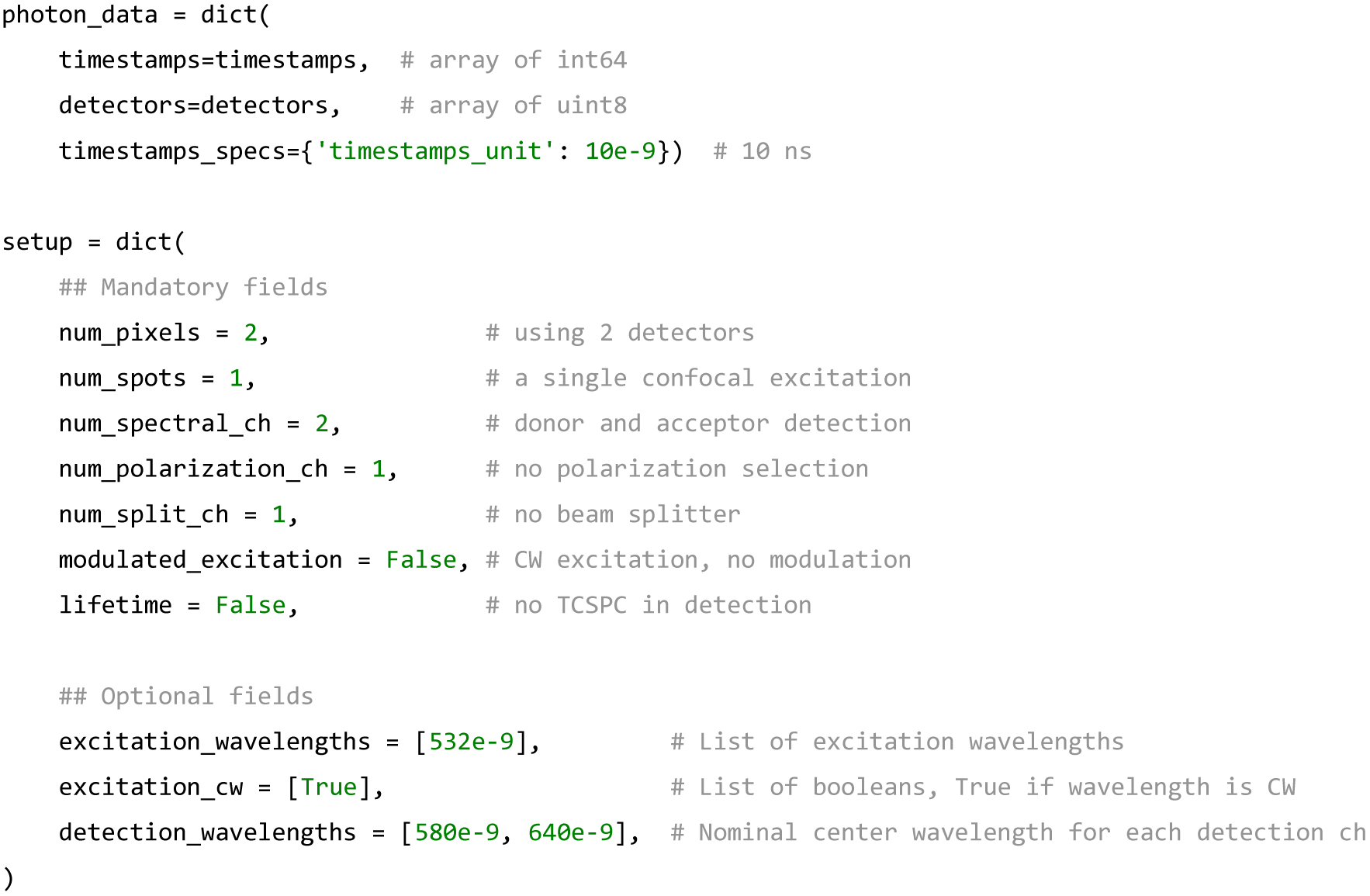

Note that a subgroup (such as *timestamps_specs* in *photon_data*) is simply a nested dictionary. We need to merge this data to create the root group of the Photon-HDF5 file, which also requires a *description* field:

~~~
data = dict(
       description = 'This is a dummy dataset which mimics smFRET data.',
       photon_data = photon_data,
       setup = setup)
~~~

Finally, the data can be saved to disk by calling the function *save_photon_hdf5* in phconvert:

~~~
import phconvert as phc
phc.hdf5.save_photon_hdf5(data, h5_fname='dummy_dataset.h5')
~~~

This will save a minimal Photon-HDF5 file. Phconvert will add a few additional fields that can be generated automatically (e.g. *acquisition_duration*, field *software* in the *identity* group, etc), a description for each field and will validate the field names and types to conform to the specifications. This file will not include a *measurement_specs* group. A complete example using non-mandatory fields (including the *measurement_specs* group) is included in phconvert and accessible at:

> http://nbviewer.ipython.org/github/Photon-HDF5/phconvert/blob/master/notebooks/Writing_Photon-HDF5files.ipynb

#### SM3.2. Saving Photon-HDF5 from non-Python programs: phforge

Users may want to implement the ability to save Photon-HDF5 files in software for data-acquisition or simulators not written in Python. As mentioned before, using phconvert would greatly simplify writing Photon-HDF5 files while ensuring compatibility with the Photon-HDF5 specifications. However, calling phconvert from other languages can be problematic. One approach is calling the Python C API to call phconvert, but this is not a particularly easy task. Alternatively, users could write Photon-HDF5 files directly using the HDF5 library for the language of choice. However, in order to ensure the creation of valid Photon-HDF5 files, the program should check the spelling of all the field names, their data types, add the official field descriptions and make sure that all the mandatory fields are presents. Most of these validations can be implemented using the Photon-HDF5 JSON specifications file. In practice, one would need to re-implement the same functionalities of phconvert, but in another language.

In order to make it easy to create validated Photon-HDF5 in any language and bypass this reimplementation step, we devised an alternative approach in which the Photon-HDF5 is created using a script called phforge(4). The process involves three simple steps:

1. Save the mandatory photon-data arrays (timestamps, detectors, nanotimes, etc…) in a plain HDF5 file.
2. Write the remaining metadata in a simple text file (in YAML format, see below).
3. Call *phforge* providing metadata and photon-data file names as input arguments to create a valid Photon-HDF5 file using phconvert.

These three steps are now briefly discussed.

1. To save the photon-data arrays, the user needs to call the HDF5 library for the language he or she chooses to use. In MATLAB for example, *timestamps* and *detectors* arrays can be saved with the following commands:

~~~
h5create('photon_data.h5', '/timestamps', size(timestamps), 'Datatype', 'int64')
h5write('photon_data.h5', '/timestamps', timestamps)
h5create('photon_data.h5', '/detectors', size(detectors), 'Datatype', 'uint8')
h5write('photon_data.h5', '/detectors', detectors)
~~~

A similar example using LabVIEW is provided at https://github.com/Photon-HDF5/photon-hdf5-labview-write.

2. The metadata file is a text-based representation of the full Photon-HDF5 structure, excluding the photon-data arrays and a few other fields automatically filled by phconvert. To store this metadata, we use YAML markup (a superset of JSON) for its simplicity and ability to describe hierarchical structures. For example, a minimal metadata file containing only mandatory fields looks as follows:
description: This is a dummy dataset which mimics smFRET data.

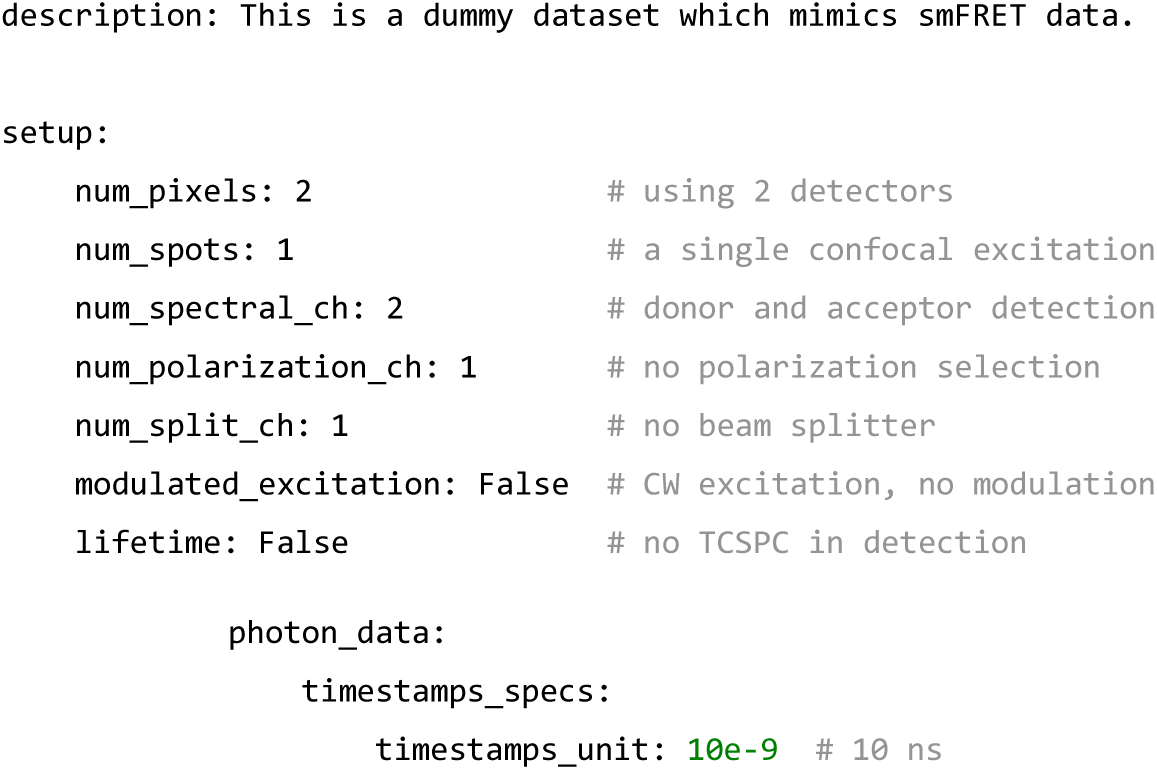

3. Finally, once metadata and photon-data files have been saved, a Photon-HDF5 file can be created calling the phforge script as follows:

~~~
phforge metadata.yaml photon-data-arrays.h5 photon-hdf5-output.hdf5
~~~

where *metadata.yaml* designates the path to the metadata file, *photon-data-array.h5* the path to the temporary HDF5 file (containing only photon-data arrays) and *photon-hdf5-output.hdf5*, the path of the destination file.

Note that the file generated with this minimal metadata example will not contain a *measurement_specs* group, which is in general necessary for a user to analyze the data. The fields corresponding to the measurement_specs group (or any other valid Photon-HDF5 field) can be added to the metadata file, so that phforge incorporates them in the final photon-hdf5 file.

The phforge script is available online(4) and easy to install on all the major operating systems. More examples of metadata files, including non-mandatory fields and measurement_specs group for different types of measurements, are available at:

> https://github.com/Photon-HDF5/phforge/tree/master/example_data

A complete example of how to create Photon-HDF5 files in MATLAB is provided at:

> https://github.com/Photon-HDF5/photon-hdf5-matlab-write

A complete example of how to create Photon-HDF5 files in LabVIEW is provided at:

> https://github.com/Photon-HDF5/photon-hdf5-labview-write

No matter which language the acquisition software is written in, it should be remembered that Photon-HDF5 files need, in most cases, some pre-processing of the data. For example, timestamps and detectors arrays needs to be separated (they are often packed in a common structure) and overflow corrections needs to be applied to timestamps (which are usually recorded with 32 or less bits by the acquisition hardware). Additionally, for μs-ALEX or ns-ALEX experiments, information on the alternation periods should be provided (although it is not mandatory). This processing can be performed either in real-time during the acquisition or in a second post-acquisition step. The latter approach naturally fits with using the phforge script for creating the final Photon-HDF5 file.

As a final note, if the developer decides to modify his or her acquisition software to save Photon-HDF5 files, he or she should also consider providing the user a simple way to input the required metadata. When converting files with a phconvert notebook, this metadata is entered using the notebook interface. When using the phforge script in custom acquisition software this metadata can be acquired by manually editing a pre-formatted YAML file or through a custom GUI which generates the metadata file automatically. The latter approach, for example, is followed in the previously linked LabVIEW example for writing Photon-HDF5 files.

### SM4. Benchmarks

#### SM4.1 Overview

In this section, we report the results of two simple read- and write-speed benchmarks for two different data files: one based on a HT3 file (created by the MicroTime 200 hardware manufactured by PicoQuant) and one based on a SPC file (created by a SPC-630 board manufactured by Becker & Hickl). The benchmarks were coded in Python 2.7 and pytables 3.1 and run on a Windows 7 x64 desktop PC with Intel^®^ Core™ i5-3570K CPU, 16GB of RAM and a 7,200 rpm SATA hard drive. The quoted values are the result of a single execution after a system reboot (to eliminate perturbation from file-system cache).

It should be noted that the value reported are only indicative and cannot be generalized because compression performances (size and speed) can significantly vary from file to file. Furthermore HDF5 allows changing one parameter, the array chunk size (the size of an on-disk-contiguous chunk of data), which can affect (in particular, improve) performance. Here, we used the default value as set by the pytables library.

We benchmarked different compression levels ranging from 0 (no compression) to 9 (maximum compression) and 2 compression algorithms: zlib (HDF5 default) and blosc (a high-speed alternative compressor developed by the pytables team and not shipped by default with the HDF5 library). It is important to note that, when using languages other than Python, using blosc requires compiling the blosc sources, which can represents a significant obstacle for the user. Therefore, we discourage the use of the blosc compressor for any file which needs to be shared. On the other hand, we included it in the benchmarks due to its impressive performance. For instance, in critical, real-time applications where the hard drive write-speed is a bottleneck (i.e. acquiring from large SPAD arrays), the blosc compressor could be used and the conversion to zlib postponed to a second, post-processing step.

#### SM4.2 Results

Read speeds were relatively uniform at all compression levels when using zlib, with values around 40 million photons per second (MP/s). In all examples, the read speed from compressed Photon-HDF5 format (compression level of 5) was around twice faster than reading from the native formats (the latter have an overhead due to byte decoding and rollover correction). It is worth noting that the read speed of the native formats depends on the specific decoding implementation, and C routines could potentially result in faster read speed (although we believe our implementation is reasonably fast).

Write speeds, on the contrary, were strongly affected by the compression level (5-10 MP/s using zlib5). Obviously, the fastest was to write uncompressed data (> 100 MP/s). Remarkably, blosc exhibited write speeds which were comparable to the uncompressed one, while significantly reducing the file size. For single-spot excitation data files, the write speed was more than enough to save data in real time even when using the slow zlib compression. For setup using large detector arrays, it would be advisable to not use any compression or only blosc compression (and instead convert files with zlib compression post acquisition or for sharing).

**Figure SM-4:**
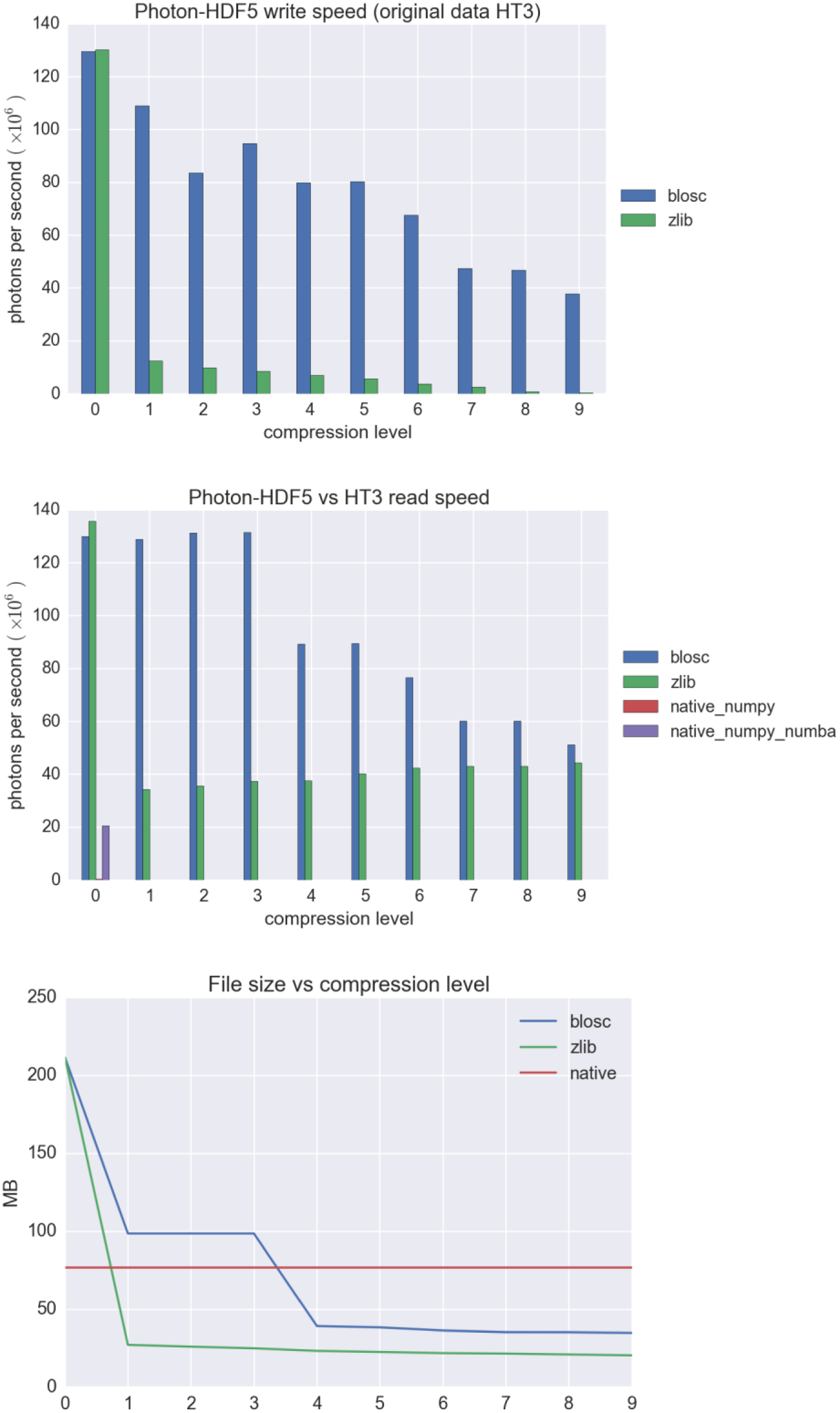
Write and Read speed benchmark using a PicoQuant HT3 input file. In the first two graphs (Write and Read speed, respectively), larger values are better, while in the last (size), smaller is better. Compression level of 0 indicates no compression.

**Figure SM-5:**
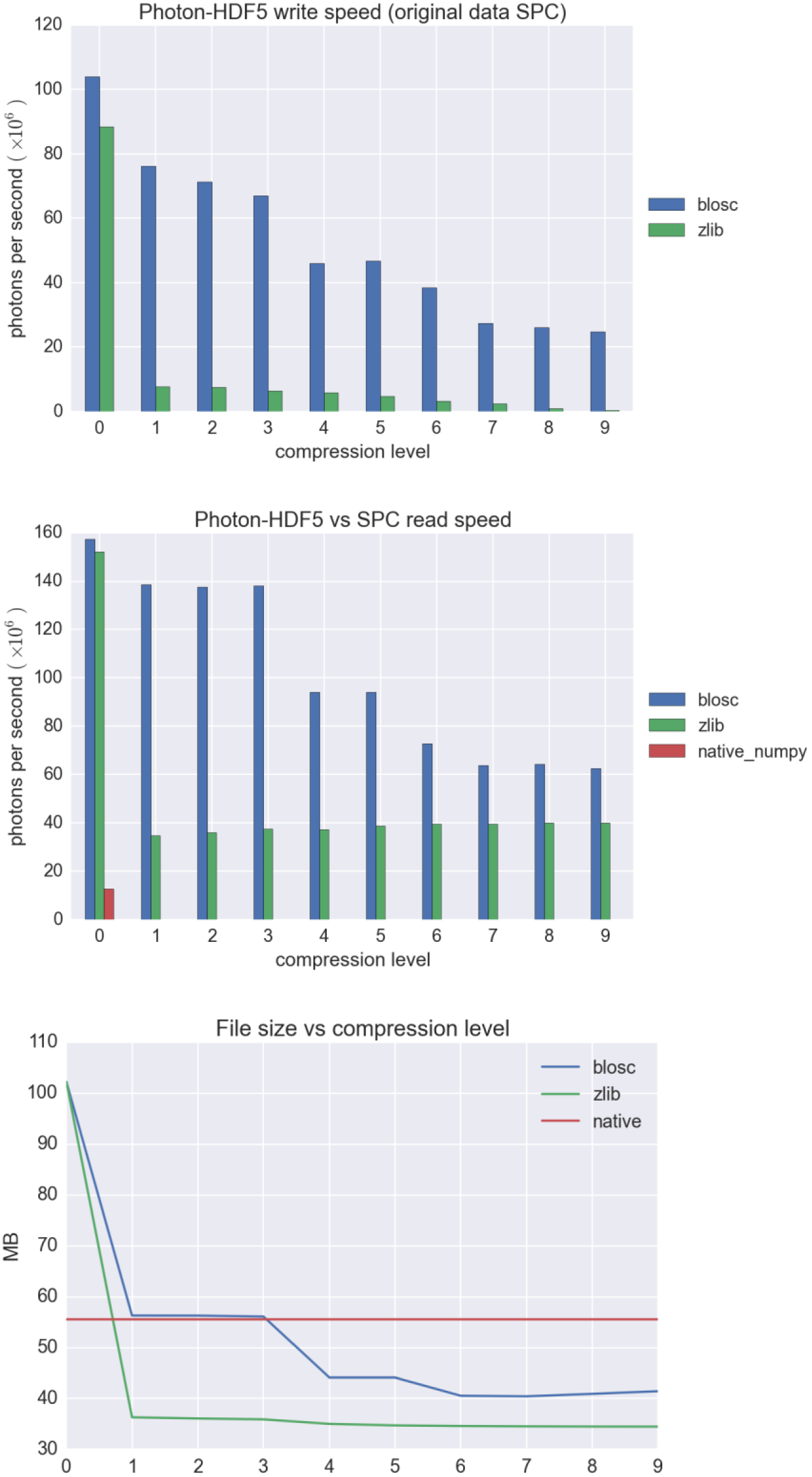
Write and Read speed benchmark using a Becker & Hickl SPC input file. In the first two graphs (Write and Read speed, respectively), larger values are better, while in the last (size), smaller is better. Compression level of 0 indicates no compression.

## Supporting Material References

1. Photon-HDF5 format definition. http://photon-hdf5.readthedocs.org/en/latest/phdata.html.
2. Reading Photon-HDF5 in multiple languages. http://photon-hdf5.github.io/photon_hdf5_reading_examples/.
3. Photon-HDF5 Reference: Reading Photon-HDF5 files. http://photon-hdf5.readthedocs.org/en/latest/reading.html.
4. phforge: a script for creating Photon-HDF5 files. http://photon-hdf5.github.io/phforge/.

